# A population-scale isoform atlas enables isoform-based trait association and regulatory analysis in lettuce

**DOI:** 10.64898/2026.11.07.765046

**Authors:** Guangbao Luo, Li Xiao, Xiaoyan Zhang, Guizhen Lan, Xiaosa Xu, Jiongjiong Chen, Hanhui Kang, Xin Wang

**Affiliations:** National Key Laboratory for Germplasm Innovation & Utilization of Horticultural Crops; Hubei Hongshan Laboratory, College of Horticulture and Forestry Sciences, Huazhong Agricultural University, 430070 Wuhan, China; Department of Plant Biology, University of California, Davis, CA 95616, USA

## Abstract

Genomic variation shapes phenotypic diversity, yet how genetic variation contributes to transcript isoform diversity remains poorly understood at the population scale. Here, we integrated transcriptomic data from 240 lettuce accessions with a lettuce pangenome to construct a comprehensive pan-isoform atlas comprising 531,657 isoforms from 49,729 transcribed loci. Isoform presence–absence variation (iPAV) captured extensive transcript diversification during domestication and horticultural differentiation, providing additional resolution beyond locus-level analyses, particularly within loci shared across the population. To facilitate the systematic analysis of isoform–trait associations, we developed ISONET-GWAS, a computational framework that integrates isoform-based association mapping, functional annotation, and network-based candidate prioritization. ISONET-GWAS recovered the known chlorophyll regulator *LsGLK*, identified 455 metabolite–isoform associations across 77 metabolites, and prioritized candidate regulators of chlorophyll metabolism supported by functional validation. Genome-wide splicing quantitative trait locus (sQTL) mapping identified 10,498 sQTL associations, revealing widespread distal genetic associations with alternative splicing and candidate trans-sQTL hotspots. Notably, we identified a trans-sQTL hotspot overlapping *LsPPR*, which encodes a pentatricopeptide repeat RNA-binding protein. Perturbation of *LsPPR* resulted in widespread changes in alternative splicing of floweing-related genes, accompanied by delayed bolting and flowering and altered splicing of flowering-related genes. Together, our study establishes a population-scale pan-isoform resource and a broadly applicable analytical strategy for isoform-level trait association, candidate prioritization, and complementary analysis of the genetic regulation of alternative splicing in crops.

## Introduction

Genetic variation contributes to phenotypic diversity through changes in gene expression, transcript processing and protein function. Although genomic variation has been extensively characterized at the DNA level, its functional consequences are implemented through the transcriptome and proteome. Transcript isoforms constitute a critical intermediate layer linking genotype to phenotype by diversifying transcript structure, expression and protein function.

Alternative splicing is a major source of transcript isoform diversity in eukaryotes, enabling a single gene to generate multiple transcript isoforms with distinct regulatory or functional properties. In plants, alternative splicing plays fundamental roles in development^1, 2^, stress responses^3, 4^, flowering time^5, 6, 7^, metabolism^8, 9^ and environmental adaptation^7, 10, 11^. Increasing evidence indicates that transcript isoform diversity is extensively shaped by natural genetic variation, including SNPs, InDels and structural variants located within splice sites, gene body or regulatory regions. For example, a 12-bp intronic deletion in DRO2 triggers intron retention and enhances drought resistance in potato^12^, while intronic retrotransposon insertions in maize WAXY alleles induce aberrant alternative splicing and exon skipping, despite their excision from host introns^13^. Genome-wide alternative splicing quantitative trait locus (sQTL) analyses in rice revealed that both SNPs and larger structural variations (SVs) extensively modulate alternative splicing via *cis*- and *trans*-acting genetic associations^9, 14^. Together, these findings illustrate how genetic variation and post-transcriptional regulation contribute to transcript isoform diversity and phenotypic variation. However, despite its biological importance, transcript isoform diversity remains much less explored than genomic variation at the population scale.

Recent advances in crop pangenomes and population genetics have expanded our understanding of genomic diversity by uncovering extensive gene presence-absence variation (PAV) and structural variation across populations^15, 16, 17^. Genome-wide association studies (GWAS) combined with expression quantitative trait locus (eQTL) analyses approach in plants^9, 14, 18, 19, 20, 21^, have successfully linked DNA polymorphisms, gene expression, and gene presence–absence variation with phenotypic diversity. PAV of *HPS* was shown to regulate seed luster in soybean^22^, PAV of three FLOWERING LOCUS C genes affecting flowering time and ecotype differentiation of *Brassica napus*^15^ and SV in tomato *WR13* regulates lipid synthesis^23^. However, these population-scale analyses remain predominantly gene-centric. Pangenomes characterize gene presence-absence variation, whereas eQTL and GWAS generally treat genes as primary functional unit. This framework overlooks transcript isoform diversity generated through alternative splicing, transcript truncation, untranslated region remodeling, alternative promoter usage and SV-associated transcript remodeling. These isoform variation frequently modifies protein domains or regulatory activity without altering gene presence or overall gene expression, suggesting that a substantial proportion of functional variation resides below the gene level. Despite the increasing availability of population transcriptomes, a comprehensive population-scale framework for characterizing transcript isoform diversity, its variation with complex traits, and integrating it with genomic variation is still lacking^24, 25^.

Lettuce (*Lactuca* sativa) is one of the world’s most widely cultivated vegetable and an important dietary source of vitamins, antioxidants and micronutrient, making it an excellent system for investigating transcript diversification during crop evolution. A single domestication event from its wild progenitor *Lactuca serriola* gave rise to the diversification of horticulture types, including crisphead, butterhead, romaine, stem, oilseed and looseleaf^26^. Previous genomic, transcriptomic and QTL cloning studies have revealed that transcript-level regulation is an important mechanism underlying lettuce domestication and agronomic trait variation^26^. For example, insertion of a CACTA transposable element into *LsKN1* creates a novel transcript that promotes leafy-head development, whereas alternative splicing of *LsGLK* attenuates chlorophyll biosynthesis and results in pale-green leaves^27, 28^ ^29^. These findings demonstrated that transcript isoforms represent an important source of functional variation during lettuce domestication and horticultural diversification, highlighting the importance of systematically characterizing transcript isoform diversity at the population level.

Here, we established a population-scale pan-isoform atlas by integrating transcriptomic data with a high-quality lettuce pangenome to systematically characterize transcript isoform diversity during domestication and horticultural diversification. Our results indicate that iPAV, particularly variation among isoforms derived from core loci, provides additional resolution for characterizing horticultural differentiation beyond locus-level variation. To enable systematic trait association and candidate prioritization at the transcript isoform level, we developed ISONET-GWAS, an isoform-based association framework integrating trait association mapping, co-functional network reconstruction and AI-assisted network interpretation. Application of this framework to pigment and metabolic traits identified trait-associated isoforms and their potential regulatory networks, including functionally validated regulators of chlorophyll accumulation. Furthermore, genome-wide sQTL mapping identified widespread distal genetic associations with alternative splicing and candidate regulatory hotspots, including regions overlapping *RBP45* and *LsPPR*. Functional characterization of LsPPR provided experimental support for its involvement in alternative splicing and flowering-time regulation, highlighting a potential relationship between transcript processing and developmental variation. Together, our study provides a population-scale isoform resource and an analytical framework for exploring isoform-level variation, trait associations, and genetic regulation, expanding the utility of population transcriptomics for functional genomics and crop research.

## Results

### An expanded genomic reference supports a population-scale lettuce pan-isoform atlas

To capture a broader range of isoform diversity in Lactuca species, we first constructed an expanded lettuce genomic reference using the representative assembled genome from stem lettuce, the wild progenitor *L. serriola*, the widely cultivated lettuce cultivar Tizian, together with the Salinas V11 reference genome. As taking the cultivar lettuce Salinas (Version 11) as reference genome, we incorporated 141,609,895 bp of non-reference sequences, which comprised 15,644 contigs from stem genome, 11,645 contigs from Tizian and 45,339 contigs from *L. serriola*. This integrative framework provided an expanded genomic foundation beyond the single reference genome, enabling a more comprehensive recovery of population-level transcript diversity. The cleaned RNA-seq reads from 240 accessions, representing multiple horticultural types and wild relatives, were aligned to this pangenome. In total, we identified 531,657 transcript isoforms derived from 49,729 transcribed loci (**Supplementary Table 1).** As transcriptome diversity is primarily driven by mRNA alternative splicing rather than the emergence of novel gene content, we collectively defined these 531,657 transcripts as the lettuce pan-isoform atlas, of which 92.1% (489,645/531,657) are multiple exons transcript. To further characterize protein-coding isoforms within the atlas, we used TransDecoder to predict protein sequences from all 531,657 isoforms and performed similarity searches against the Swiss-Prot and UniProt databases, identifying 265,653 protein-coding isoforms derived from 21,362 loci. Notably, approximately three-quarters of these loci contributed two or more isoforms, highlighting the extensive prevalence of isoform diversification at the population scale (**Supplementary Table 1**). The atlas included 23,819 isoforms associated with 16,070 loci absent from the previous lettuce reference annotation, including 1,342 putative protein-coding isoforms from 930 newly annotated loci (**Supplementary Table 1; Supplementary Fig. 1a,b**). Functional annotation of the newly identified coding isoforms included categories related to plant–pathogen interactions and receptor-mediated signaling (**Supplementary Fig. 1c-e; Supplementary Table 2**).

We next classified isoforms by their detection frequency across the lettuce population. A total of 9,386 isoforms were detected in every accession (core), 7,514 in 90–99% of accessions (soft-core), and 60,787 in 5–90% of accessions (dispensable); 453,970 isoforms (approximately 85%) were detected in fewer than 5% of accessions and classified as private (**Figure 1a, b**).

**Figure 1.**
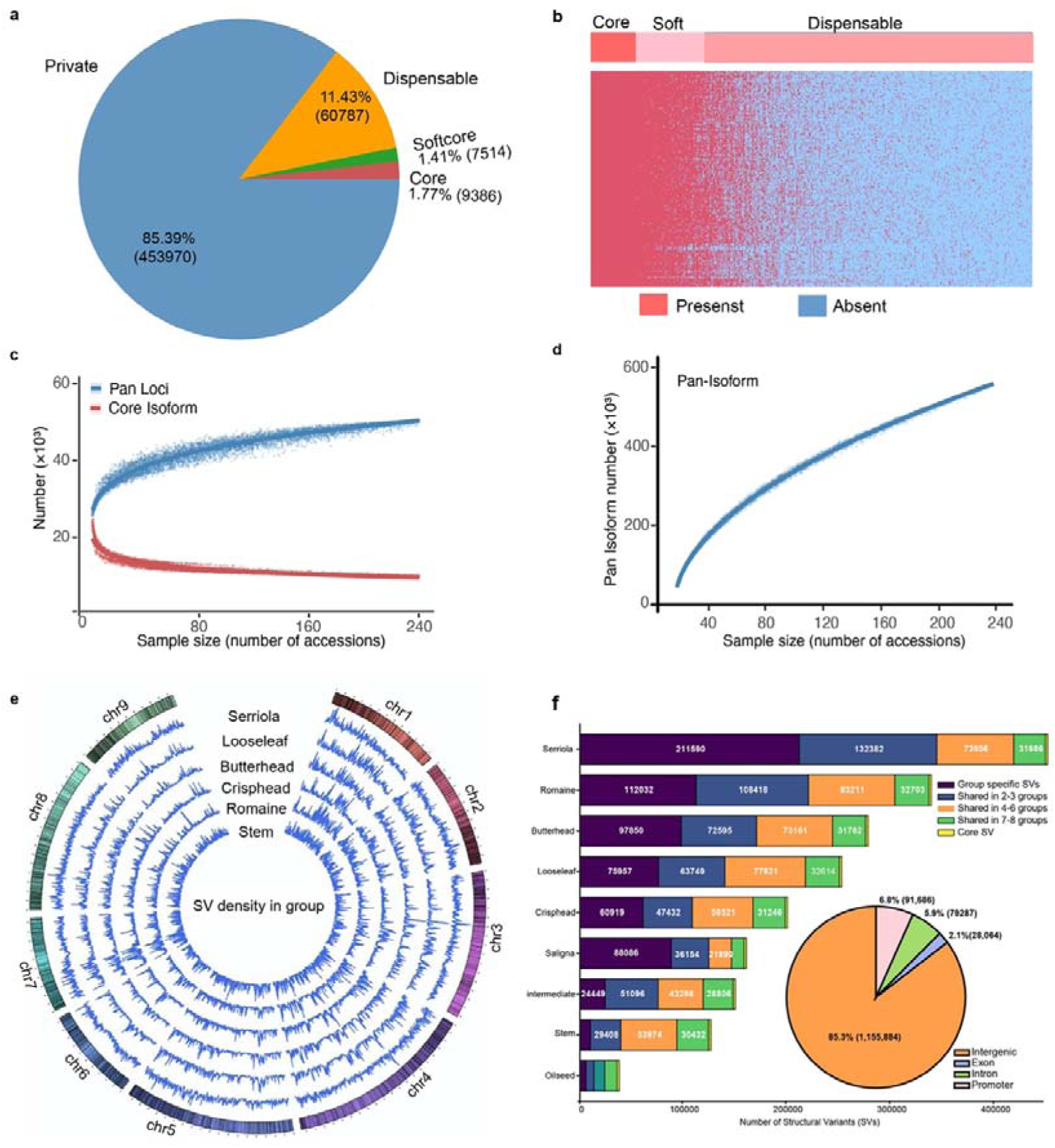
Population-scale pan-isoform discovery and genomic structural variation in lettuce. **a,** Proportions and numbers of transcript isoforms classified as core, soft-core, dispensable and private according to their presence frequencies across 240 lettuce accessions. **b**, Presence/absence variation of core, soft-core and dispensable isoforms across the 240 accessions. Red and blue indicate isoform presence and absence, respectively, with isoform categories indicated at the top. **c**, Accumulation curves showing changes in the numbers of pan-transcript loci (blue) and core isoforms (red) with increasing accession number. **d,** Accumulation curve showing the number of non-redundant pan-isoforms identified with increasing population size, based on randomized combinations of the 240 accessions. **e,** Genome-wide distribution of SV density of lettuce groups. The SV densities were anchored along each chromosome. **f,** Distribution of group-specific, shared and core SVs across lettuce groups. The inset pie chart shows the genomic distribution of high-confidence SVs in intergenic, promoter, intronic and exonic regions.

The predominance of low-frequency isoforms highlights the extensive diversity of the lettuce pan-isoform repertoire, with most identified isoforms detected in only a small fraction of the population. This pattern is consistent with substantial transcript-level diversity across lettuce accessions, although the observed detection frequencies may also be influenced by differences in transcript abundance and sequencing-based detection. Rarefaction analysis based on iterative random sampling of accessions suggested that the number of pan-transcript loci approached saturation, while the number of core isoforms stabilized at a finite level (**Figure 1c**). In contrast, the total number of detected isoforms continued to increase with population size without reaching an apparent plateau (**Figure 1d**). Collectively, these results indicate that isoform diversity remains less saturated than locus-level diversity, highlighting the additional transcriptomic variation captured at the isoform level.

### Isoform-level variation provides additional resolution beyond locus-level PAV

To explore the relationship between isoform presence–absence variation (iPAV) and nearby genomic variants, we analyzed whole-genome resequencing data from 129 representative lettuce accessions to construct a population-scale map of structural variants (SVs) and insertions/deletions (InDels). A total of 1,354,881 SVs (≥ 50 bp) and 4,170,600 InDels (1-49 bp, MAF ≥ 0.05) in the lettuce resequencing datasets were found using the PSVGT framework^30^. Classification of SVs by horticultural groups showed that most detected SVs were shared across multiple cultivated types and wild accessions, with only a small proportion of core SVs, suggesting that these SVs originated before the diversification of cultivated lettuce (**Figure 1e, f**). Annotation of high-confidence SVs and InDels revealed that these variants frequently overlapped promoter, intronic and exonic regions, with 6.8% located in 3-kb promoters, 5.9% in introns, and 2.1% in exons. We used Fisher exact test to evaluate associations between *cis* variants and isoforms. The result showed that *cis* insertion/deletion were associated with the 7,339 isoforms (**Figure 2a, b**), with approximately 45% of the associated variants located within gene bodies (**Figure 2c**). However, 94% of isoforms were not significantly associated with *cis* variants, due to lower frequency or no variant found around the gene. These results indicate substantial population-level variation in isoform presence/absence that is not explained simply by genomic PAV, and reflect differences in transcript isoform generation or retention between wild and cultivated lettuce.

**Figure 2.**
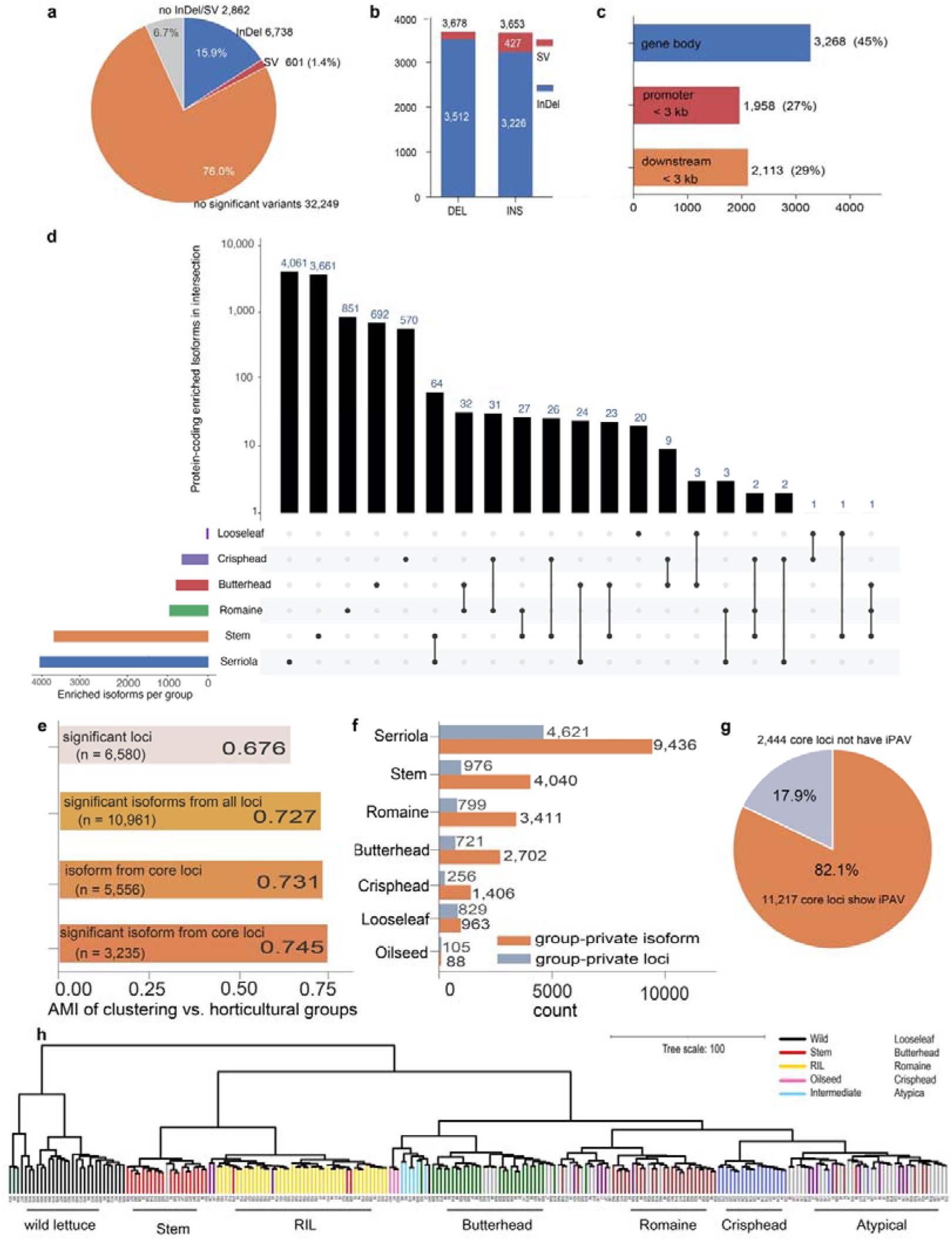
Isoform-level variation provides additional resolution for lettuce horticultural differentiation. **a-c,** *cis* variants types and location that associated iPAV. d, The enriched isoform among each lettuce groups. **e,** AMI between hierarchical clustering of the population and the horticultural groups, computed on Fisher-significant group-enriched loci or isoform (FDR < 0.05): significant loci, significant isoforms from all loci, isoforms from core loci and significant isoforms from core loci. **f,** Group-private loci versus group-private isoforms (detected in ≥2 accessions) for each horticultural type. **g,** Of the 13,661 core loci detected in all accessions, 82.1% of the core loci exhibited iPAV. **h,** The genome-wide intron-splicing clustering of lettuce population, VaSP was used to calculate intron alternative value.

To investigate whether isoform-level measurements distinguish domestication status and horticultural groups more clearly than locus-level PAV, we compared iPAV frequencies among different horticultural groups and identified 26,964 isoforms associated with domestication and horticultural diversification, including 11,993 isoforms uniquely enriched in a single group (**Figure 2d; Supplementary Fig. 2; Supplementary Table 3**). The strongest differentiation was observed between cultivated lettuce and its wild progenitor, *L. serriola*, with 3,239 isoforms significantly enriched in cultivated lettuce and 4,154 enriched in *L. serriola*, with many of these isoforms annotated to metabolic functions (**Figure 2d; Supplementary Fig. 2; Supplementary Table 3a**). Among cultivated horticultural types, stem lettuce displayed the most extensive isoform differentiation, consistent with our previous observation that stem lettuce harbored the highest number of detected transcripts, further supporting an expanded isoform repertoire associated with stem lettuce improvement. The KEGG annotation of the 3,239 protein-coding isoforms revealed that terms related to spliceosome, plant-pathogen interaction and motor proteins were more favorable among cultivars than *L. serriola* (**Supplementary Fig. 3**). Compared with wild groups, metabolic pathways were highly enriched, suggesting that cultivated lettuce possesses a great repertoire of transcripts involved in the metabolic processes. To further assess the contribution of isoform variation to horticultural differentiation, we compared the ability to distinguish horticultural groups of isoform- and locus-level PAV. Hierarchical clustering of all horticulture lettuces accessions based on isoforms resolved horticultural types more effectively than locus level, with adjusted mutual information (AMI) values of 0.73 and 0.68, respectively (**Figure 2e**). Notably, 29.5% of group-enriched isoforms originated from core loci shared across all analyzed accessions, and these isoforms achieved the highest clustering, achieving an AMI of 0.75 (**Figure 2e**). Moreover, 22,046 group-private isoform events supported by at least two accessions were identified, compared 8,307 group-specific loci (**Figure 2f**). Of the 13,661 core loci detected in all accessions, 82.1% of the loci exhibited isoform PAV specific enriched in distinct type of lettuce (**Figure 2g**). Together, these results indicate that isoform-level variation, particularly variation among isoforms within core loci, provides greater resolution than locus-level PAV for distinguishing lettuce horticultural groups and characterizing their differentiation.

To assess splicing variation independently of isoform present-absent variation, we also quantified 36,054 alternative splicing events using VaSP across the lettuce population. The clusters of alternative splicing events broadly reflected horticultural differentiation, although looseleaf accessions were distributed among several branches (**Figure 2h**). These results provide independent evidence that population structure is also reflected in quantitative splicing variation, further supporting a pervasive association between transcript isoform variation and population diversification.

### ISONET-GWAS integrates isoform–trait association mapping with network-based candidate prioritization

The extensive isoform-level variation captured by the pan-isoform atlas provides an opportunity to develop approaches that use transcript isoforms as molecular features for trait analysis and candidate prioritization. We therefore developed ISONET-GWAS, which uses iPAV as a molecular association feature and combines the resulting trait associations with functional annotation and network candidate prioritization (**Figure 3a**). To evaluate the performance of ISONET-GWAS, we first applied this framework to the chlorophyll a+b profile from metabolism data collected in previous study ^28, 29^. ISONET-GWAS identified the *LsGLK* isoform PAV as the most significantly associated isoform for chlorophyll a+b, with -log_10_ (P) =15.80 (**Figure 3b,c; Supplementary Table 4**). In comparison, a previously reported SNP-based GWAS^29^ detected an association within a 5.6-Mb genomic interval containing *LsGLK*, with a peak −log₁₀(P) value of 6.60. The associated isoforms were also observed in metabolites carotenoids, chlorophyll b and total pigment, demonstrating that ISONET-GWAS could provide complementary resolution for identifying associated loci at the isoform level relative to conventional SNP-based GWAS. In addition to *LsGLK*, ISONET-GWAS identified several isoforms associated with chlorophyll-related traits (**Supplementary Fig. 4a; Supplementary Table 5**). Among the strongest pigment-associated isoforms, FERONIA (*TCONS_00248141*), involved in photorespiratory flow and photosystem maintenance^31, 32^, and a myosin-2-like isoform (*TCONS_00119940*), involved in chloroplast photorelocation^33, 34^, co-associated with the *LsGLK* allele multiple pigment traits. Additional candidates included a chlorophyll a/b-binding protein (*TCONS_00359288*), LIGHT-DEPENDENT SHORT HYPOCOTYLS3 (*TCONS_00470920*), and the MSL2-like isoform *TCONS_00349681*, implicated in chloroplast growth and division^35, 36^ (**Supplementary Table 5**). Together, these results indicate that ISONET-GWAS identifies trait-associated isoforms with plausible functional links to the relevant biological pathways, providing candidates for further investigation of the relationships between genetic variation, transcript diversity and metabolic traits.

**Figure 3.**
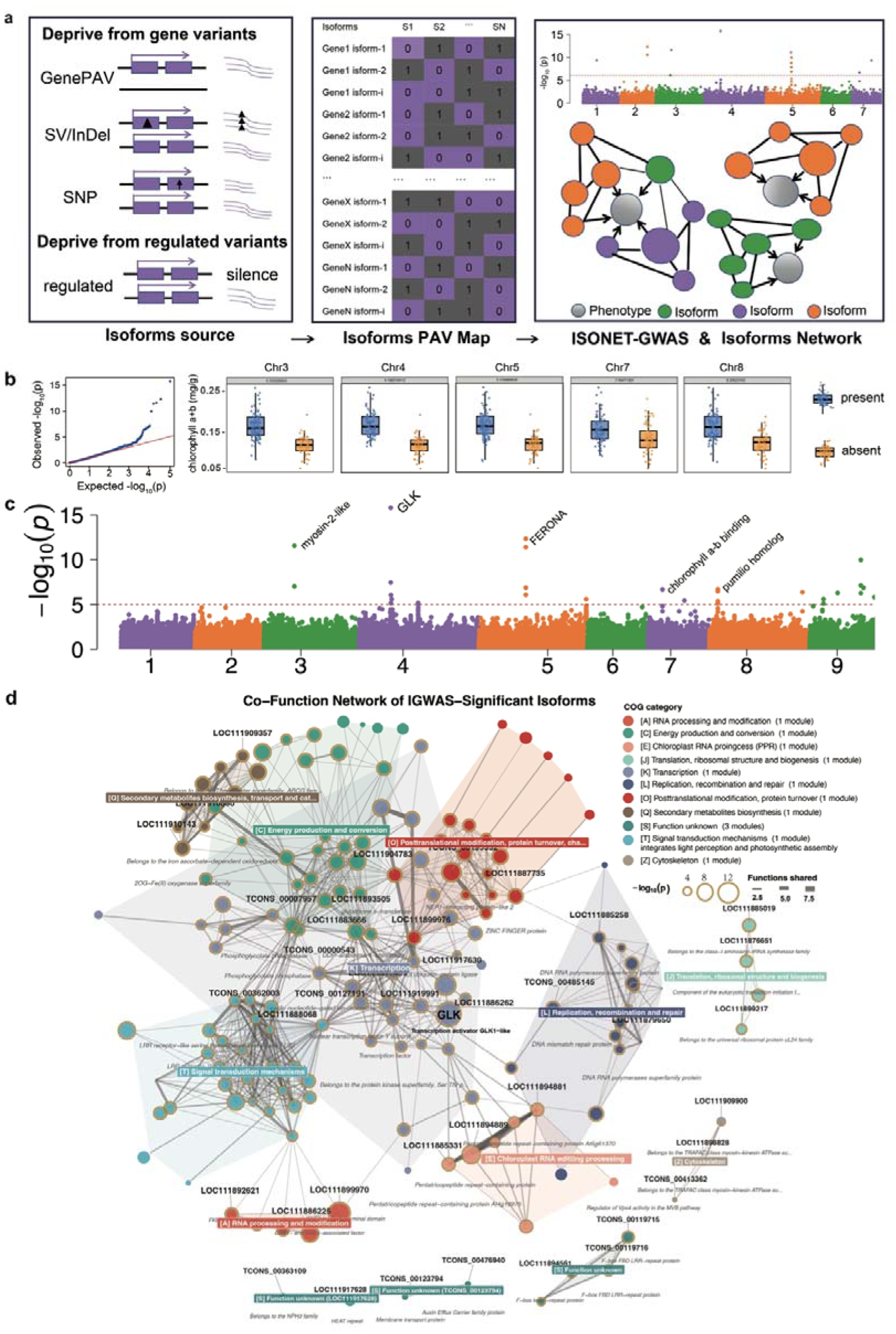
ISONET-GWAS identifies chlorophyll-associated isoforms and reconstructs functional networks. **a**, Schematic overview of the ISONET-GWAS framework. **b**, Quantile-quantile plot of ISONET-GWAS associations with chlorophyll a+b content and box plots showing differences in chlorophyll a+b content between accessions with (blue) and without (orange) selected associated isoforms. **c,** Manhattan plot of associated isoforms. The x-axis represents the genomic location of each isoform, and y-axis represents -log10 of the association P value. **d**, Co-functional network of isoforms significantly associated with chlorophyll a+b content.

Biological regulatory pathways are often highly complex, involving multiple functional layers and numerous interconnected components, making it challenging to systematically interpret trait-associated isoform networks. Moreover, association strength alone does not necessarily reflect regulatory importance, making it difficult to identify the modules and relationships most relevant to a given trait. To address this interpretive bottleneck, ISONET-GWAS combined co-functional network reconstruction based on shared GO terms, PFAM domains, and KEGG pathways with AI-assisted regulatory interpretation to integrate functional evidence and co-expression-based network expansion. In chlorophyll metabolism, the resulting network connected GLK1-like transcriptional regulators, chlorophyll-binding proteins, metabolic signaling components, and chloroplast/RNA-processing PPR proteins, with repeated AI-assisted interpretations at different time points consistently suggesting coordinated transcriptional, post-transcriptional, and metabolic regulation (**Figure 3d; Supplementary Table 6**). Within the PPR associated module, *LOC111886547* was prioritized as a candidate hub. Knockout plants displayed pale-green, speckled leaves and altered chlorophyll accumulation (**Supplementary Fig. 5**), supporting a role for this candidate in chlorophyll-related phenotypes. These findings provide experimental support for the biological relevance of the prioritized candidate and suggest that combining ISONET-GWAS with AI-assisted network interpretation could facilitate the identification of candidate regulators from complex isoform-associated networks.

### ISONET-GWAS reveals isoform–metabolite associations and candidate regulatory networks in lettuce

To evaluate the applicability of ISONET-GWAS across diverse quantitative traits, we extended the analysis to 77 metabolites and identified 455 significant metabolite–isoform associations (**Supplementary Table 7**). Among the associated isoforms, most encode proteins with functions directly related to their corresponding metabolites, supporting the biological relevance of the identified associations. For example, glutamate was significantly associated with *TCONS_00170824*, which encodes a 2-oxoglutarate-dependent dioxygenase (2ODD). Members of the 2ODD family require 2-oxoglutarate (2OG) as an essential co-substrate, which is directly derived from glutamate metabolism via transamination or deamination^37, 38, 39, 40^, linking glutamate availability to downstream 2OG-dependent processes. In addition, histidine was associated with *TCONS_00361595*, a PfkB-family carbohydrate kinase involved in generating ribose-5-phosphate^41, 42^, the direct precursor for phosphoribosyl pyrophosphate (PRPP) required in histidine biosynthesis^43, 44, 45^. Glucose-6-phosphate was associated with *LOC111885767*, encoding phosphoenolpyruvate carboxykinase, a key regulatory enzyme in gluconeogenesis that determines the production of glucose-6-phosphate from non-carbohydrate precursors^46, 47, 48, 49^.

To further explore the regulatory architecture underlying metabolic diversity, we extended the network analysis to all 77 metabolites. These networks revealed recurrent functional modules involving signaling proteins, metabolic enzymes, transcription factors and ubiquitin-mediated protein turnover (**Supplementary Table 7**). For example, the proline-associated network was enriched in receptor-like kinases and NB-ARC/LRR proteins, suggesting a potential link between stress signaling and proline accumulation, whereas the fucose network contained enzymes involved in glycosylation and sugar metabolism. Transcription factors, including bZIP, bHLH and MYB family members, together with ubiquitin-related proteins, were also recurrently represented across multiple metabolic networks, highlighting the complexity of metabolite regulation. Notably, *TCONS_00532049* and *TCONS_00265730* were associated with 47 and 24 metabolites, respectively, suggesting potential candidate hubs associated with multiple metabolic traits. Together, these results demonstrate the applicability of ISONET-GWAS to diverse metabolic traits and its potential to identify associated isoforms and network modules, providing isoform-level genetic resources for dissecting metabolic regulation and facilitating precision breeding of favorable metabolic traits in lettuce.

### Complementary sQTL analysis reveals widespread genetic associations with alternative splicing and candidate regulatory hotspots

To complement the isoform–trait association analyses, we performed genome-wide sQTL mapping to investigate the genetic basis of alternative splicing diversity and its association with genomic variation in lettuce. Genome-wide sQTL mapping using 3S score value generated by VaSP as the phenotype identified 10,498 sQTLs for the 4,206 alternatively spliced introns in 2,665 genes (**Figure 4a; Supplementary Table 8**). The sQTL distribution showed that only 12.4% of sQTLs were *cis*-sQTL, whereas 87.6% were *trans*-sQTL (**Figure 4a, b**). Among 139 genes with trait-associated isoforms and testable sQTL phenotypes, 32 exhibited significant sQTL associations, including 21 with trans-only associations, 10 with both cis- and trans-associations, and one with cis-only associations. The remaining 107 genes showed no significant sQTL association (**Supplementary Table 9**). These findings indicate that isoform–trait associations and sQTL mapping capture partially overlapping but distinct aspects of transcript variation, highlighting their complementary potential for investigating the genetic regulation and phenotypic relevance of transcript isoforms.

**Figure 4.**
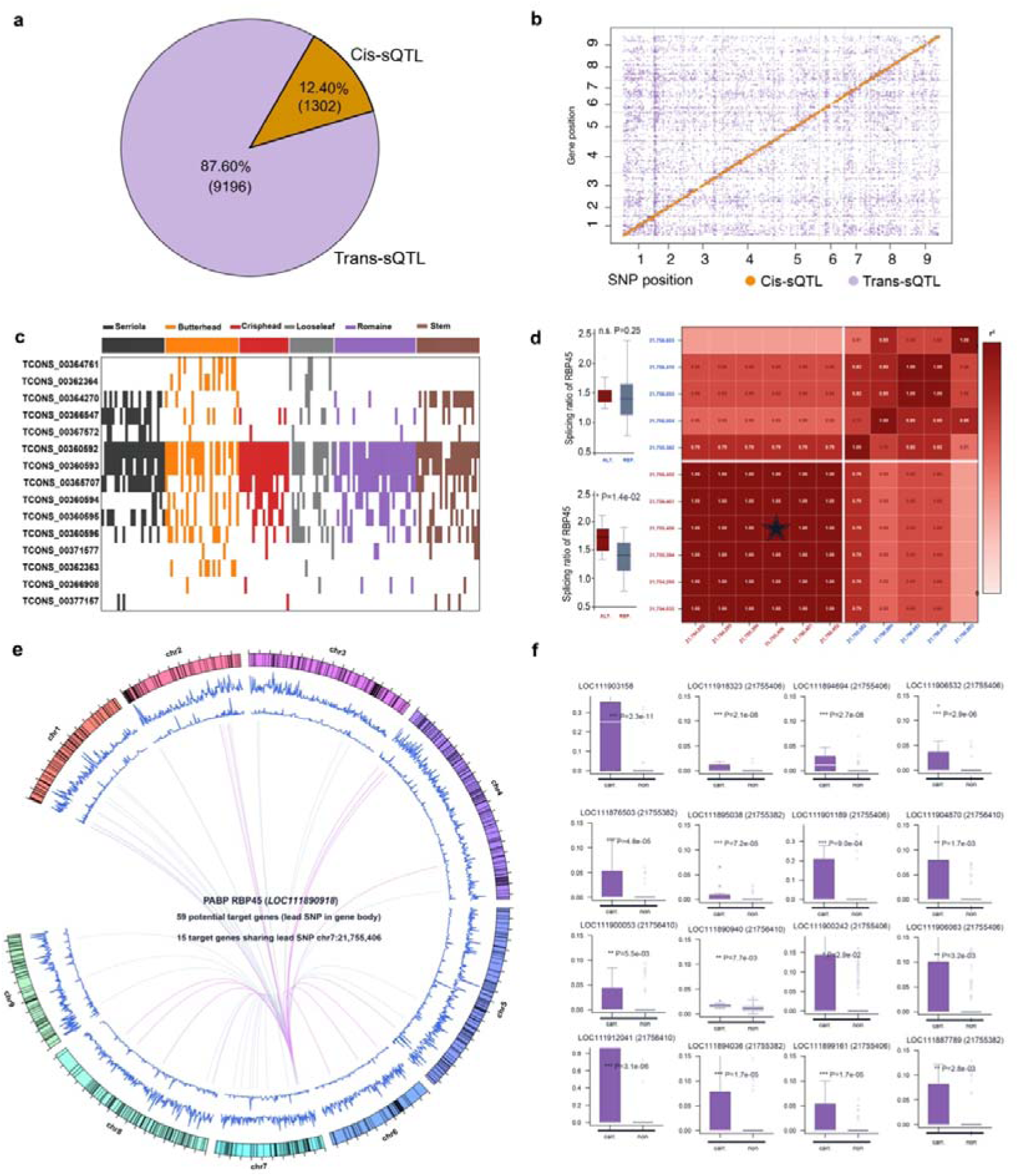
Genome-wide sQTL landscape and RBP45 splicing regulation in lettuce. **a-b,** Overview of sQTL components and their genomic distribution. **c,** Distribution of RBP45 transcript isoforms across lettuce groups. **d,** Linkage disequilibrium (LD; (r^2)) heatmap of 11 linked variants (9 SNPs and 2 InDels) in two LD blocks, separated by white lines: block 1 (red), comprising six SNPs, and block 2 (blue), comprising five variants. Each cell represents the pairwise (r^2) value (white-to-red color scale, 0–1), and the asterisk indicates the *cis*-associated SNP at chr7:21,755,406. Box plots show RBP45 splicing ratios associated with different alleles in the two LD blocks. **e,** Circos plot showing potential RBP45 target genes whose lead sQTL SNPs map within the RBP45 gene body. Purple chords represent the 15 target genes sharing the same *cis* lead SNP at chr7:21,755,406, whereas light-grey chords represent the 44 target genes associated with other lead SNPs. **f,** Box plots showing the splicing ratios of target genes in accessions carrying different alleles at RBP45-associated loci. “non” and “carr” denote the reference and alternative alleles of RBP45, respectively.

To determine whether trans-sQTLs exhibit hotspot patterns similar to those observed for eQTLs, we examined their genome-wide distribution and identified genomic regions with a significant enrichment of trans-sQTLs. A total of 564 candidate hotspots, as well as the potential master splicer within each hotspot were identified, associated with splicing of 1,875 genes (**Supplementary Table 10-12**). One hotspot overlapped the splicing-factor gene RBP45 and was associated with alternative splicing in 59 genes (**Supplementary Table 11**). Nine SNPs and two InDels within the RBP45 gene body were associated with variation in RBP45 isoform usage and represent candidate local variants for further investigation (**Figure 4c**). For example, the RBP45 isoform *TCONS_00364761* was distinctly enriched in butterhead, while *TCONS_00364270* enriched in both *L. serriola* and stem lettuce (**Figure 4c**). Among the 59 target genes whose alternative splicing was associated with this sQTL hotspot, 15 shared the lead variant at chr7:21,755,406 (**Figure 4e**). This genomic block was also associated with alternative splicing of both downstream target genes and RBP45 itself, suggesting a link between cis-regulatory variation in *RBP45* and trans-acting splicing regulation (**Figure 4e, f**). Together, these findings reveal a genetic architecture of alternative splicing in lettuce characterized by widespread trans-sQTL associations and the clustering of distal splicing signals at candidate regulatory hotspots.

### Functional characterization of LsPPR links a candidate trans-sQTL hotspot to alternative splicing and flowering-time variation

Beyond splicer regulation gene located within hotspot, we also identified a major hotspot that located a single gene--*LsPPR,* which encodes a pentatricopeptide repeat (PPR) RNA-binding protein (**Figure 5a**). The lead SNPs associated with isoforms of 71 target genes were located within the gene body of *LsPPR,* suggesting that this locus represents a candidate master splicer of alternative splicing in lettuce (**Supplementary Table 12**). To further investigate the regulatory function of *LsPPR*, we generated *LsPPR* knockout lines by CRISPR-Cas9 in lettuce inbred line E26J. Despite screening across multiple generations, we did not obtain viable homozygous *LsPPR* knockout mutants or high-expression overexpression lines. These observations suggest that strong perturbation of *LsPPR* may be associated with impaired development or reduced viability and are consistent with the possibility that *LsPPR* dosage is important for normal development. We therefore selected heterozygous knockout and moderate overexpression lines for RNA-seq analysis to investigate the potential role of *LsPPR* in alternative splicing.

**Figure 5.**
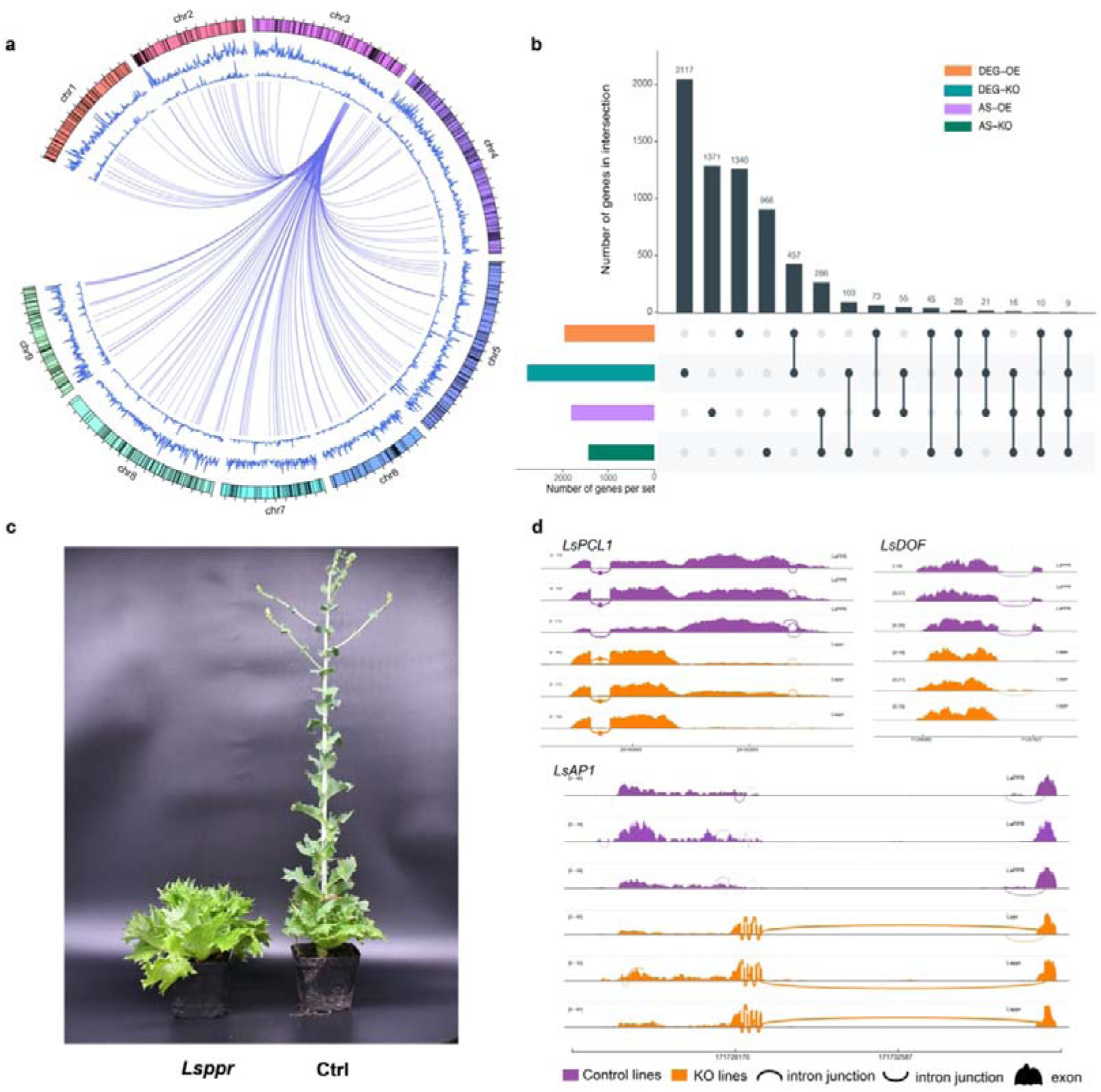
*LsPPR* represents a trans-sQTL hotspot associated with alternative splicing and flowering-time regulation. **a,** Circos plot showing the genome-wide distribution of sQTLs (black lines). The first blue track represents gene density in 1-Mb windows, and the second blue track represents the density of genes associated with sQTLs in 500-kb windows. Purple chords connect the *LsPPR* hotspot to its putative target genes. **b,**UpSet plot showing the overlap between genes with altered alternative splicing in *LsPPR* overexpression (OE) and knockout (KO) lines. **c,** Delayed bolting and flowering of *Lsppr* mutants compared with control plants. **d,** Sashimi plots showing RNA-seq read coverage and splice-junction arcs for the flowering-time and circadian-rhythm genes *LsAP1*, *LsDOF* and *LsPCL1* in control and *Lsppr* mutant plants. *LsAP1* shows a shift from predominantly multi-exon-skipping isoforms in control plants to predominantly full-length transcripts in the mutants, whereas *LsDOF* and *LsPCL1* exhibit altered first-exon usage.

RNA-seq analysis of *LsPPR* heterozygous mutant revealed widespread alterations in alternative splicing, with 1,408 genes showing significant alternative splicing changes relative to control plants (**Figure 5b; Supplementary Tables 12-13**). Among these, eight genes overlapped with the 71 predicted *trans*-sQTL target genes, including AP-1 complex subunit (*LOC111887177*), disease resistant protein (*LOC111908479*), ribonucleo protein (*LOC111903831*), protein kinase (*LOC111891467*) and long non-coding RNA (*LOC111878742*). The limited overlap may reflect differences between natural genetic variation captured by population-based sQTL mapping and the broader splicing responses induced by experimental perturbation of *LsPPR* in a single genetic background, potentially compounded by the partial loss of function in the heterozygous mutant. Similarly, RNA-seq analysis of overexpression line identified 1,802 genes with altered AS patterns, including eight genes overlapping with the predicted *trans*-sQTL target set (**Supplementary Table 12 & 14**). These genes included WD40 (*LOC111882684*) whose homolog regulates embryo development in *Arabidopsis*. In addition, disease resistant protein (*LOC111908479*), ribonucleo protein (*LOC111903831*), protein kinase (*LOC111891467*) were also shown altered splicing in overexpression lines. The overlap between genes with experimentally altered splicing and the genetically predicted target set provides partial support for the predicted regulatory relationships. The widespread splicing changes observed in both the *LsPPR* heterozygous knockout and overexpression lines further suggest that *LsPPR* is involved in the regulation of alternative splicing at the genome-wide level.

Consistent with its extensive effects on transcript processing, *Lsppr* heterozygous mutants exhibited significantly delayed in bolting and flowering compared with control plants (**Figure 5c**). The differential alternative splicing events in the mutants were enriched in genes that involved in flowering-related pathways, a potential association between altered transcript processing and the flowering phenotype (**Supplementary Table 15**). For instance, MADS-box gene *LsAP1*(*LOC111888924*) participating in circadian rhythm and vernalization response, multi-exons skipping isoform were highly enriched in the control groups while in the heterozygous mutant groups the complete transcripts were predominate (**Figure 5d**). Another MADS-box gene *LsFBP24* displayed abundant isoforms with intron retention in control plants, while such transcripts were largely absent in the mutant (**Supplementary Fig. 6**). In addition, the DOF zinc finger *LsDOF* (*LOC111903145*), a negative regulator of photoperiodism and flowering, showed altered usage of first exon between mutants and control plants (**Figure 5d**). Similarly, the circadian rhythm response regulator *LsPCL1* (*LOC111907478*) also displayed alternative of first exon usage (**Figure 5d**). Collectively, these findings support a role for *LsPPR* in alternative splicing and flowering-time regulation in lettuce, with altered splicing of flowering-related genes accompanying delayed bolting and flowering in heterozygous mutants. Although the specific splicing events underlying the flowering phenotype remain to be determined, these results highlight a potential connection between transcript isoform regulation and genomic variation in lettuce.

## Discussion

Population genomics has transformed our understanding of crop evolution by revealing extensive genomic variation across natural population. However, gene-level analyses do not capture all the ultimate functional units through which genetic variation shaped phenotypes. Instead, the functional consequences of genomic variation are largely mediated through transcript isoform, which represents an important yet underexplored layer of functional variation in crop evolution. Despite this central role, population-scale characterization of transcript isoforms diversity remains much less developed than population genomics in plants. However, several representative examples have demonstrated that changes in either transcript isoforms abundance or structure can produce striking phenotypic consequences in lettuce^27, 29, 50^. Our population-scale pan-isoform atlas provides a transcript-level resource for systematically characterizing variation that is not fully represented by conventional gene-level analyses. The enrichment of distinct isoforms among horticultural groups suggests that domestication and diversification have been accompanied by extensive changes in transcript usage, highlighting isoform diversification as a potentially important component of lettuce evolution.

A major contribution of this study is the development of ISONET-GWAS, an isoform-based association and network analysis framework that extends population-level trait analysis beyond conventional genomic variant approaches. By combining iPAV association mapping with functional network reconstruction, co-expression analysis, and candidate prioritization, ISONET-GWAS provides a systematic workflow for exploring trait-associated transcript diversity. Conventional GWAS associates phenotypic variation with genomic polymorphisms, whereas the causal regulatory consequences often remain difficult to interpret because multiple variants converge on the same regulatory outcome. In contrast, ISONET-GWAS directly employs transcript isoform PAV as association markers, establishing a more immediate link between molecular regulatory and corresponding phenotype. Consequently, ISONET-GWAS recovered the known chlorophyll regulator *LsGLK* and identified functionally related isoforms associated with the same biological pathway (**Figure 3; Supplementary Fig. 5**). Instead of identifying isolated candidate genes within the large LD region, ISONET-GWAS provides a systems framework for connecting phenotypes to functional transcript isoforms, particularly for regulatory pathways underlying complex trait. Notably, the incorporation of AI-assisted interpretation further facilitates the functional annotation, organization, and biological interpretation of complex networks, enabling the identification of potential regulatory modules underlying metabolic traits. The successful functional validation of a predicted network hub involved in chlorophyll regulation further demonstrated the capability of this approach to uncover biologically meaningful regulators from complex multi-layer networks (**Supplementary Fig. 5**).

Accumulating evidence suggests that genomic variants influence transcript diversity and that transcript isoforms contributed substantially to phenotypic variation^27, 29, 50, 51, 52, 53^. However, these relationships have largely been investigated independently. By integrating isoform analysis, genomic variation and genome-wide sQTL mapping, we established a comprehensive analytical framework for investigating the relationships between genomic variation and transcript isoform diversity, providing a foundation for exploring their potential contributions to phenotypic variation. In the present study, ISONET-GWAS directly associates transcript isoforms with metabolic traits, whereas sQTL analysis identifies the genomic variants associated with quantitative differences in alternative splicing. Among 139 genes with associated isoforms and testable sQTL phenotypes, only 32 exhibited significant sQTL associations. This partial overlap may reflect differences in the molecular features captured by the two approaches, as iPAV characterizes transcript detection patterns, whereas sQTL analysis focuses on quantitative variation in alternative splicing. Thus, these two approaches provide complementary insights into transcript variation rather than necessarily identifying identical sets of genes. Notably, the phenotype in such a framework is not limited to agronomic traits but can also include molecular phenotypes, such as gene expression levels. For example, the expression level of a key regulator or metabolic enzyme could itself be treated as a quantitative trait, enabling the identification of isoforms associated with variation in its expression. Together, ISONET-GWAS and sQTL mapping provide complementary analytical perspectives for utilizing population-scale isoform variation in functional genomics, supporting trait-associated candidate prioritization and the identification of putative splicing regulatory loci.

*PPR* genes show considerable evolutionary diversity across plant species, although the extent of variation differs among individual genes. Previous studies have also reported natural sequence variation in some *PPR* genes, particularly at residues involved in RNA recognition^54^. Such variation may affect RNA-binding specificity and the processing of different RNA targets. In lettuce, whether natural *LsPPR* alleles differ in their RNA targets and regulatory effects remains unknown. Moreover, as many PPR proteins function in organellar RNA processing, it is unclear whether the altered splicing of flowering-related genes observed in *LsPPR* mutants reflects direct or indirect regulation.

Although this study focuses on lettuce, the framework presented here is broadly applicable to other crop species as population transcriptomic resources continue to expand. Future incorporation of long-read transcript sequencing, single-cell transcriptomics, spatial transcriptomics and environmental perturbation datasets will enable increasingly complete characterization of population pan-isoforms and their regulatory dynamics across tissues and developmental stages. Similar to how pangenomes have reshaped our understanding of genome diversity, we envision population-scale pan-isoforms becoming a fundamental resource for dissecting functional variation beyond the gene level. Together, the pan-isoform atlas and ISONET-GWAS framework offer resources and methods for investigating transcript-level variation and prioritizing trait-associated candidates in lettuce, with potential applications to other crop populations.

## Materials and Methods

### Construction of lettuce pangenome

The representative genomes/contigs were downloaded from public databases, including those from the Salinas, stem lettuce, Tizian and *L. serriola* (TKI-340)^55, 56^. To identify non-reference genomic sequences, assembled contigs were aligned to the lettuce reference genome (V11) using QUAST^57^ with the parameters: threads 10, --min-identity 90, and --min-alignment 300. The unaligned contigs and sequences longer than 500 bp were retained, and redundant sequences were removed using CD-HIT^58^ with the parameters -T 48 -c 0.9 -d 0 -M. To minimize potential contamination, sequences whose best hits fell outside the green plant lineage, or that mapped to known plant mitochondrial or chloroplast genomes, were identified as potential contaminants and discarded. The final non-redundant sequences were combined with the Salinas V11 reference genome to construct the lettuce pangenome for subsequent population-scale transcriptome analyses.

### Pan-isoform construction

The lettuce pan-isoform atlas was constructed using population-scale RNA-seq data aligned to the expanded lettuce genomic reference. The raw RNA-seq reads were processed using fastp (v0.23.2) program to remove adapter sequence and low-quality reads with the default parameters^59^. The clean reads were subsequently aligned to the constructed pangenome using STAR (v2.7.10a) program^60^. The known and novel transcripts were assembled and quantified by the StringTie (version 2.2.1) with default parameters^61^. The resulting transcript annotation were compared using gffcompare (v0.12.6) program^62^ to investigate isoform absence and presence among the population. In parallel, genomic variants were identified and annotated using the PSVGT (v1.0) framework^30^.

### Identification of horticultural type specific enriched isoforms

To identify isoforms associated with horticultural differentiation of lettuce, we compared the frequency of isoforms across different horticulture types. For each isoform, a two-tailed Fisher’s exact test was performed to assess differences in its presence frequency between a given horticultural type and all remaining types combined. The resulting P-values were adjusted for multiple testing using the Benjamini–Hochberg procedure, and a false discovery rate (FDR) threshold of 0.05 was applied to determine statistical significance. Isoforms meeting this criterion and exhibiting a higher frequency in a particular horticultural type were designated as horticultural type specific enriched isoforms.

### Comparison of isoform- and locus-level resolution of horticultural groups

To compare the resolution of isoform- and locus-level presence–absence variation (PAV) in horticultural differentiation, group enriched isoforms and loci were identified using two-sided Fisher’s exact tests as described above. The pairwise Jaccard distances were calculated from binary PAV matrices, followed by average-linkage hierarchical clustering into seven clusters. The clustering performance was evaluated using adjusted mutual information (AMI) between the resulting clusters and horticultural group classifications. AMI was calculated separately for significant loci, significant isoforms from all loci, and enriched isoforms derived from core loci shared by the all horticultural accessions. Group-private loci and isoforms were defined as features detected exclusively within a single horticultural group, with group-private isoforms required to be present in at least two accessions to exclude singletons.

### ISONET-GWAS and co-functional-expression network reconstruction

ISONET-GWAS was developed to integrate isoform-trait association mapping with trait-anchored regulatory network reconstruction. For association mapping, isoform presence/absence variation was encoded as binary genetic marker map and analyzed using two complementary analyses. Analysis I included isoform PAV markers from 240 RNA-sequenced accessions, whereas Analysis II additionally incorporated gene-body structural variants in the subset of accessions with both transcriptomic and genomic data. The metabolic traits were obtained from previous study^28, 29^, and subjected to rank-based inverse-normal transformation. The association testing was performed using EMMAX^63^ with a Balding–Nichols kinship matrix and principal components as covariates. Isoform–trait associations exceeding −log₁₀(P) > 5.5 were considered significant and the significant isoform–trait pair across the two analyses was retained. Candidate isoforms were annotated through EggNOG database^64^.

Trait-anchored regulatory networks were reconstructed through two complementary layers. In Layer 1 (co-func), significant isoforms were taken as nodes to connect between each other based on GO terms, PFAM domains or KEGG pathways after removal of broad or generic terms. Network modules were identified using community detection implemented in the R package igraph^65^. Each module was then characterized and ranked by an agentic AI component (DeepSeek V4 Pro) using a plant-metabolomics-specific system prompt. Based on GO, Pfam, KEGG and COG annotations, the AI agent summarized the predominant biological functions of each module, assigned an interpretable module name, generated mechanistic hypotheses linking module members to the associated trait with a confidence level (high/medium/low), and ranked modules according to predicted trait relevance. Based on functional annotations and trait information, DeepSeek was used to help select high-confidence modules for subsequent co-expression analysis. This step was optional and did not affect the GWAS analysis or the construction of the co-function network. To assess the consistency of AI-assisted interpretation, we repeated the analysis at different time points using the same functional annotations and trait information. Although some variation in the generated descriptions was observed, the main biological interpretations remained consistent across runs.

In Layer 2 (co-func-exp), high-confidence, trait-relevant modules were expanded by incorporating isoforms sharing the functional terms, allowing one expansion step to 500 isoforms per module. A co-expression network was then constructed using the isoform-level TPM values within the expanded module (Spearman correlation r ≥ 0.65, FDR ≤ 0.05, minimum TPM 0.1). The network centrality was evaluated using degree, betweenness, and module eigengene connectivity (kME). Isoforms were subsequently re-ranked using a combined score integrating ISONET-GWAS significance (30% weight) and co-expression centrality (70% weight). Isoforms or genes exhibiting high network centrality despite weak or absent association at primary ISONET-GWAS significance threshold were identified as potential “rising hubs” based on a rank rise ≥ 10, and p-value < 0.01. The resulting networks were visualized using igraph^65^.

### Alternative splicing detection and sQTL analysis

The alternative splicing events across the lettuce population was quantified using VaSP (version 1.20.0) package to calculate each alternative Introns splicing score in R program^66^. The final alternative splicing score of each intron was used as quantitative phenotypes to conduct genome wide sQTL mapping using EMMAX^63^. The principal components of the population and Kinship relationship matrix was calculated by FlashPCA2^67^ and EMMAX-KIN^63^, respectively. The LD among significant SNPs was calculated using PLINK (version 1.9) with parameters --r2 --ld-window-r2 0 --ld-window 9999999 --ld-window-kb 10000000. Adjacent significant SNPs within physical distance of 4.7Mb and exhibiting strong LD (r^2^ > 0.1) were merged into the sQTL blocks for subsequent analysis.

### sQTL hotspot analysis

To identify genomic regions enriched in splicing QTL (sQTLs), The hot_scan was used to analyze the hotspot of sQTL with default parameters^68^. The significant sQTLs were first grouped into independent blocks based on linkage disequilibrium, and the lead SNP with the lowest association P-value in each block was selected for hotspot scanning, with alternative splicing events provided as the corresponding splicing patterns. The genome was then scanned using a sliding-window approach, and the number of lead SNPs within each window was compared with a null distribution generated through random permutation of lead SNP positions across the genome. Windows containing significantly more lead SNPs at empirical false discovery rate < 0.05 were defined as sQTL hotspots. Within each hotspot, the SNP associated with the largest number of target genes was designated as the master sQTL.

### Generation of transgenic lettuce lines and transcriptome analysis

The CRISPR/Cas9 tetra-gRNA PL2B knockout vector was constructed. Target sites were designed with CRISPR-P v2.0 (http://crispr.hzau.edu.cn/CRISPR2/), replacing the 20-nt N-stretch in the forward primer with the gRNA sequence and the reverse primer with its reverse complement. Pair forward and reversed guide RNA with adapters were used to amplify tRNA templates (**Supplementary Table 16**). The purified PCR products were assembled into BsaI-HF-digested PL2B backbone via homologous recombination (OKclon Kit, Accurate Biology, Hunan, China For overexpression, the coding sequence of *LsPPR*, excluding the stop codon, was amplified from E26J cDNA and inserted into the BsaI-HF-linearized plGFP vector under the control of the LsUBI promoter. The resulting constructs were introduced into Escherichia coli DH5α and subsequently transformed into Agrobacterium tumefaciens GV3101 after sequence verification.

Lettuce (*Lactuca sativa* cv. E26J) was used as the genetic background for generating LsPPR overexpression (*LsPPR-OE*) and CRISPR/Cas9 knockout (*LsPPR-KO*) lines. All the plants were cultivated in a controlled-environment growth room under a 16-h light/8-h dark photoperiod. The temperature was maintained at 22 ± 1 °C, relative humidity at 65–70%. Young, fully expanded leaves (the third to fifth true leaves from the apex) were collected from two-month-old wild-type, *LsPPR*-OE and *LsPPR*-KO plants. Three biological replicates were collected for each genotype for RNA-seq libraries construction, with each replicate comprising pooled leaf tissues from three independent plants.

All sequencing reads were processed using fastp to remove adapter sequences and low-quality reads^59^. The clean reads were then aligned to the reference genome using HISAT2 with default parameters^69^. Differential expression analysis was conducted using R package DESeq2^70^, and absolute log2 fold change > 1 and adjusted P value < 0.05 were defined as differentially expressed genes. The intron alternative splicing was assessed using VaSP^66^. The identified intron splicing events were compared between groups, and events with a fold change > 1.5 and P value < 0.05 were regarded as significantly changed intron alternative splicing events.

## Acknowledgements

We thank the high-performance computing platform at the National Key Laboratory of Crop Genetic Improvement at Huazhong Agricultural University. This research was supported by grants from the National Key Research and Development Program of China (2023YFF1000100), the National Natural Science Foundation of China award no. 32573026, the scientific research start-up funding from Hubei Hongshan Laboratory (No. 11020102), the Fundamental Research Funds for the Central Universities (No.2662024SZ005), and the Young Scientist Fostering Funds for the National Key Laboratory for Germplasm Innovation & Utilization of Horticultural Crops (Horti-PY-2023-004).

## Author contributions

X.W., J.C. and H.K. conceived and supervised this work. G.B.L. designed and implemented the pan-isoform and ISONET-GWAS framework. L.X., X.Z. G.Z.L., and J.C. performed plant transformation and RNA library preparation of transgene plants. L.X., G.B.L and X.X. implemented data analysis. G.B.L., L.X., X.W. and H.K. wrote the manuscript.

## Conflict of interest

The authors declare that they have no competing interests in relation to this work.

## Data availability

The RNA-seq data used in this study were retrieved from the NCBI BioProject database under the project PRJNA394784. The population DNA-seq data generated in this study have been deposited in the NCBI BioProject database under accession number PRJNA1545312.

## Code availability

The ISONET-GWAS framework and demo are available at https://github.com/lgbTime/ISONET-GWAS. The pan-isoform construction pipeline and custom scripts used in this study are available at https://github.com/lgbTime/Lettome_pan-isoform_ISONET-GWAS_benchmarks.

## Notes

### Competing Interest Statement

The authors have declared no competing interest.

## References

1. Zhu C-Y, et al. Alternative splicing in regulating plant development and abiotic stress response. Journal of Experimental Botany 77, 3550–3563 (2026).

2. Liu R, et al. Natural variation in OsMADS1 transcript splicing affects rice grain thickness and quality by influencing monosaccharide loading to the endosperm. Plant Communications 6, 101178 (2025).

3. Thatcher SR, et al. Genome-Wide Analysis of Alternative Splicing during Development and Drought Stress in Maize Plant Physiology 170, 586–599 (2016).

4. Innes PA, Goebl AM, Smith CCR, Rosenberger K, Kane NC. Gene expression and alternative splicing contribute to adaptive divergence of ecotypes. Heredity 132, 120–132 (2024).

5. Syed NH, Kalyna M, Marquez Y, Barta A, Brown JWS. Alternative splicing in plants &#x2013; coming of age. Trends in Plant Science 17, 616–623 (2012).

6. Li P, Tao Z, Dean C. Phenotypic evolution through variation in splicing of the noncoding RNA COOLAIR. Genes Dev 29, 696–701 (2015).

7. Hanemian M, et al. Natural variation at FLM splicing has pleiotropic effects modulating ecological strategies in Arabidopsis thaliana. Nature Communications 11, 4140 (2020).

8. Wang F, et al. An alternative splicing caused by a natural variation in BnaC02.VTE4 gene affects vitamin E and glucosinolate content in rapeseed (Brassica napus L.). Plant Biotechnol J 23, 1535–1547 (2025).

9. Zhang H, et al. Population-level exploration of alternative splicing and its unique role in controlling agronomic traits of rice. The Plant Cell 36, 4372–4387 (2024).

10. Ma S, et al. Regulation of alternative splicing in the ABA signaling pathway of plants. New Phytol 250, 115–124 (2026).

11. Wang X, et al. Discovery of a DFR gene that controls anthocyanin accumulation in the spiny Solanum group: roles of a natural promoter variant and alternative splicing. The Plant Journal 111, 1096–1109 (2022).

12. Innes PA, Kane NC, Smith CCR. Evolutionary genetics of alternative splicing in plants. New Phytologist 249, 1155–1160 (2026).

13. Varagona MJ, Purugganan M, Wessler SR. Alternative splicing induced by insertion of retrotransposons into the maize waxy gene. The Plant Cell 4, 811–820 (1992).

14. Doll NM. New insights into alternative splicing in rice using population-level transcriptomics. The Plant Cell 36, 4276–4277 (2024).

15. Song J-M, et al. Eight high-quality genomes reveal pan-genome architecture and ecotype differentiation of Brassica napus. Nature Plants 6, 34–45 (2020).

16. Zhou Y, et al. Graph pangenome captures missing heritability and empowers tomato breeding. Nature 606, 527–534 (2022).

17. Zhang Y, et al. Telomere-to-telomere Citrullus super-pangenome provides direction for watermelon breeding. Nature Genetics 56, 1750–1761 (2024).

18. Xu G, et al. The incomplete dominance of gene expression controlled by Trans-eQTL hotspots contributes to heterosis in maize. Nature Communications, (2026).

19. Lei M, et al. Comprehensive analysis of 1,771 transcriptomes from 7 tissues enhance genetic and biological interpretations of maize complex traits. G3 Genes|Genomes|Genetics 15, jkaf140 (2025).

20. Zhao J, et al. eQTL Meta-Analysis Reveals Conserved and Population-Specific Regulatory Variation Underlying Nutritional Trait Evolution and Domestication in Tomato. Advanced Science n/a, e19899 (2026).

21. Zhang L, et al. Genome-wide analysis of expression quantitative trait loci (eQTLs) reveals the regulatory architecture of gene expression variation in the storage roots of sweet potato. Horticulture Research 7, 90 (2020).

22. Liu Y, et al. Pan-Genome of Wild and Cultivated Soybeans. Cell 182, 162–176.e113 (2020).

23. Wang X, et al. Genome of Solanum pimpinellifolium provides insights into structural variants during tomato breeding. Nature communications 11, 5817 (2020).

24. Guo W, et al. A barley pan-transcriptome reveals layers of genotype-dependent transcriptional complexity. Nature Genetics 57, 441–450 (2025).

25. White B, et al. De novo annotation reveals transcriptomic complexity across the hexaploid wheat pan-genome. Nature Communications 16, 8538 (2025).

26. Zhang L, et al. RNA sequencing provides insights into the evolution of lettuce and the regulation of flavonoid biosynthesis. Nature communications 8, 2264 (2017).

27. Yu C, et al. Upregulation of a KN1 homolog by transposon insertion promotes leafy head development in lettuce. Proceedings of the National Academy of Sciences 117, 33668–33678 (2020).

28. Zhang W, et al. Dissection of the domestication-shaped genetic architecture of lettuce primary metabolism. The Plant Journal 104, 613–630 (2020).

29. Zhang L, Qian J, Han Y, Jia Y, Kuang H, Chen J. Alternative splicing triggered by the insertion of a CACTA transposon attenuates LsGLK and leads to the development of pale-green leaves in lettuce. The Plant Journal 109, 182–195 (2022).

30. Luo G, Xiao L, Fei Z, Wang X. A sensitive and accurate framework for population-scale structural variant discovery and genotyping across sequence types. bioRxiv, 2026.2001.2010.698766 (2026).

31. Jiang W, et al. FERONIA regulates salt tolerance in Arabidopsis by controlling photorespiratory flux. The Plant Cell 36, 4732–4751 (2024).

32. Shin SY, et al. FERONIA Confers Resistance to Photooxidative Stress in Arabidopsis. Frontiers in Plant Science Volume 12 - 2021, (2021).

33. Krzeszowiec W, Gabryś H. Phototropin Mediated Relocation of Myosins in Arabidopsis thaliana. Plant Signal Behav 2, 333–336 (2007).

34. Sattarzadeh A, Krahmer J, Germain AD, Hanson MR. A Myosin XI Tail Domain Homologous to the Yeast Myosin Vacuole-Binding Domain Interacts with Plastids and Stromules in *Nicotiana benthamiana*. Molecular Plant 2, 1351–1358 (2009).

35. Veley Kira M, Marshburn S, Clure Cara E, Haswell Elizabeth S. Mechanosensitive Channels Protect Plastids from Hypoosmotic Stress During Normal Plant Growth. Current Biology 22, 408–413 (2012).

36. Haswell ES, Meyerowitz EM. MscS-like Proteins Control Plastid Size and Shape in Arabidopsis thaliana. Current Biology 16, 1–11 (2006).

37. Yang X, et al. Rice Ferredoxin-Dependent Glutamate Synthase Regulates Nitrogen–Carbon Metabolomes and Is Genetically Differentiated between japonica and indica Subspecies. Molecular Plant 9, 1520–1534 (2016).

38. Temple SJ, Vance CP, Stephen Gantt J. Glutamate synthase and nitrogen assimilation. Trends in Plant Science 3, 51–56 (1998).

39. Forde BG, Lea PJ. Glutamate in plants: metabolism, regulation, and signalling. Journal of Experimental Botany 58, 2339–2358 (2007).

40. Fletcher SC, Coleman ML. Human 2-oxoglutarate-dependent oxygenases: nutrient sensors, stress responders, and disease mediators. Biochemical Society Transactions 48, 1843–1858 (2020).

41. Elkin SR, Kumar A, Price CW, Columbus L. A broad specificity nucleoside kinase from Thermoplasma acidophilum. Proteins: Structure, Function, and Bioinformatics 81, 568–582 (2013).

42. Gao B, et al. Structural analysis and functional study of phosphofructokinase B (PfkB) from Mycobacterium marinum. Biochemical and Biophysical Research Communications 579, 129–135 (2021).

43. Hove-Jensen B, Andersen Kasper R, Kilstrup M, Martinussen J, Switzer Robert L, Willemoës M. Phosphoribosyl Diphosphate (PRPP): Biosynthesis, Enzymology, Utilization, and Metabolic Significance. Microbiology and Molecular Biology Reviews 81, 10.1128/mmbr.00040-00016 (2016).

44. Hove-Jensen B. Mutation in the phosphoribosylpyrophosphate synthetase gene (prs) that results in simultaneous requirements for purine and pyrimidine nucleosides, nicotinamide nucleotide, histidine, and tryptophan in Escherichia coli. Journal of Bacteriology 170, 1148–1152 (1988).

45. Ames BN, Martin RG, Garry BJ. The First Step of Histidine Biosynthesis. Journal of Biological Chemistry 236, 2019–2026 (1961).

46. Holyoak T, Sullivan SM, Nowak T. Structural Insights into the Mechanism of PEPCK Catalysis. Biochemistry 45, 8254–8263 (2006).

47. Yang J, Kalhan SC, Hanson RW. What Is the Metabolic Role of Phosphoenolpyruvate Carboxykinase? *. Journal of Biological Chemistry 284, 27025–27029 (2009).

48. Lardy H, Hughes PE. Regulation of Gluconeogenesis at Phosphoenolpyruvate Carboxykinase. In: Current Topics in Cellular Regulation (eds DeLuca M, Lardy H, Cross RL). Academic Press (1984).

49. Yu S, Meng S, Xiang M, Ma H. Phosphoenolpyruvate carboxykinase in cell metabolism: Roles and mechanisms beyond gluconeogenesis. Molecular Metabolism 53, 101257 (2021).

50. Jia Y, et al. Up-regulation of LsKN1 promotes cytokinin and suppresses gibberellin biosynthesis to generate wavy leaves in lettuce. J Exp Bot 73, 6615–6629 (2022).

51. Tossolini I, Mencia R, Arce AL, Manavella PA. The genome awakens: transposon-mediated gene regulation. Trends in Plant Science 30, 857–871 (2025).

52. Bhati M, Mapel XM, Lloret-Villas A, Pausch H. Structural variants and short tandem repeats impact gene expression and splicing in bovine testis tissue. Genetics 225, iyad161 (2023).

53. Sousa-Luís R, Carmo-Fonseca M. Tools and tactics for studying alternative splicing. Nature Reviews Genetics 27, 642–660 (2026).

54. Barkan A, et al. A combinatorial amino acid code for RNA recognition by pentatricopeptide repeat proteins. PLoS Genet 8, e1002910 (2012).

55. Wei T, et al. Whole-genome resequencing of 445 Lactuca accessions reveals the domestication history of cultivated lettuce. Nature Genetics 53, 752–760 (2021).

56. Verwaaijen B, et al. Assembly of the Lactuca sativa, L. cv. Tizian draft genome sequence reveals differences within major resistance complex 1 as compared to the cv. Salinas reference genome. Journal of Biotechnology 267, 12–18 (2018).

57. Gurevich A, Saveliev V, Vyahhi N, Tesler G. QUAST: quality assessment tool for genome assemblies. Bioinformatics 29, 1072–1075 (2013).

58. Fu L, Niu B, Zhu Z, Wu S, Li W. CD-HIT: accelerated for clustering the next-generation sequencing data. Bioinformatics 28, 3150–3152 (2012).

59. Chen S. fastp 1.0: An ultra-fast all-round tool for FASTQ data quality control and preprocessing. iMeta 4, e70078 (2025).

60. Dobin A, et al. STAR: ultrafast universal RNA-seq aligner. Bioinformatics 29, 15–21 (2013).

61. Pertea M, Pertea GM, Antonescu CM, Chang T-C, Mendell JT, Salzberg SL. StringTie enables improved reconstruction of a transcriptome from RNA-seq reads. Nature Biotechnology 33, 290–295 (2015).

62. Pertea G, Pertea M.

63. Kang HM, et al. Variance component model to account for sample structure in genome-wide association studies. Nature Genetics 42, 348–354 (2010).

64. Huerta-Cepas J, et al. eggNOG 5.0: a hierarchical, functionally and phylogenetically annotated orthology resource based on 5090 organisms and 2502 viruses. Nucleic Acids Res 47, D309–d314 (2019).

65. Csárdi G, Nepusz T. The igraph software package for complex network research.) (2006).

66. Yu H, Du Q, Campbell M, Yu B, Walia H, Zhang C. Genome-wide discovery of natural variation in pre-mRNA splicing and prioritising causal alternative splicing to salt stress response in rice. New Phytologist 230, 1273–1287 (2021).

67. Abraham G, Qiu Y, Inouye M. FlashPCA2: principal component analysis of Biobank-scale genotype datasets. Bioinformatics 33, 2776–2778 (2017).

68. Silva IT, Rosales RA, Holanda AJ, Nussenzweig MC, Jankovic M. Identification of chromosomal translocation hotspots via scan statistics. Bioinformatics 30, 2551–2558 (2014).

69. Kim D, Paggi JM, Park C, Bennett C, Salzberg SL. Graph-based genome alignment and genotyping with HISAT2 and HISAT-genotype. Nature Biotechnology 37, 907–915 (2019).

70. Love MI, Huber W, Anders S. Moderated estimation of fold change and dispersion for RNA-seq data with DESeq2. Genome Biology 15, 550 (2014).

