## Supplementary material for "A population-scale isoform atlas enables isoform-based trait association and regulatory analysis in lettuce": Supplymentary Figure

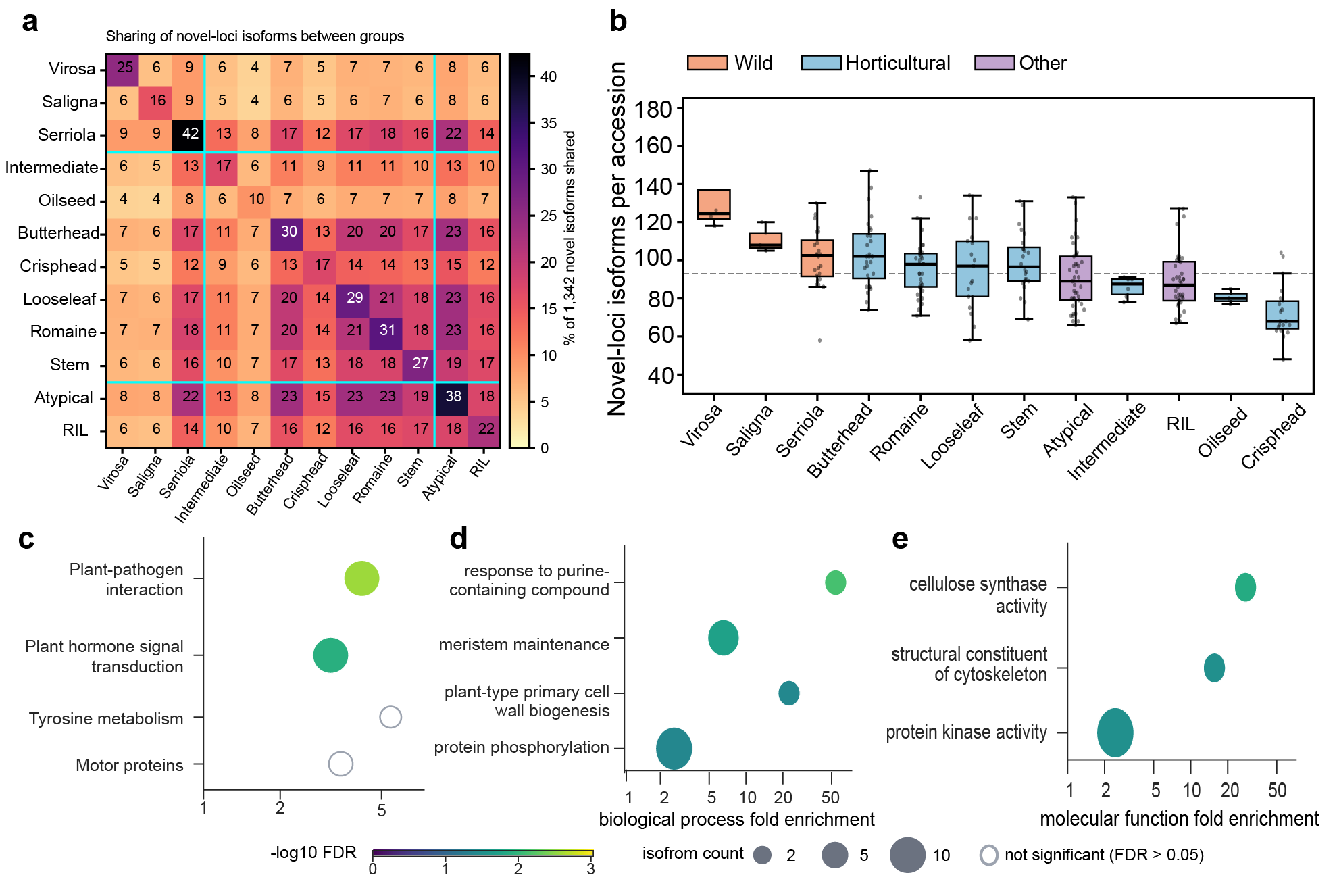

**Supplementary Figure 1. Novel-loci isoform distribution and functional enrichment analysis of protein-coding isoforms.** a, Isoform from novel-loci sharing within lettuce group. b, boxplot of the novel-loci isoform from each lettuce group. c, KEGG pathway enrichment of the protein-coding isoforms from novel-loci. d, GO biological process enrichment analysis of protein-coding isoforms from novel-loci. e, GO molecular function enrichment analysis of protein-coding isoforms from novel-loci.

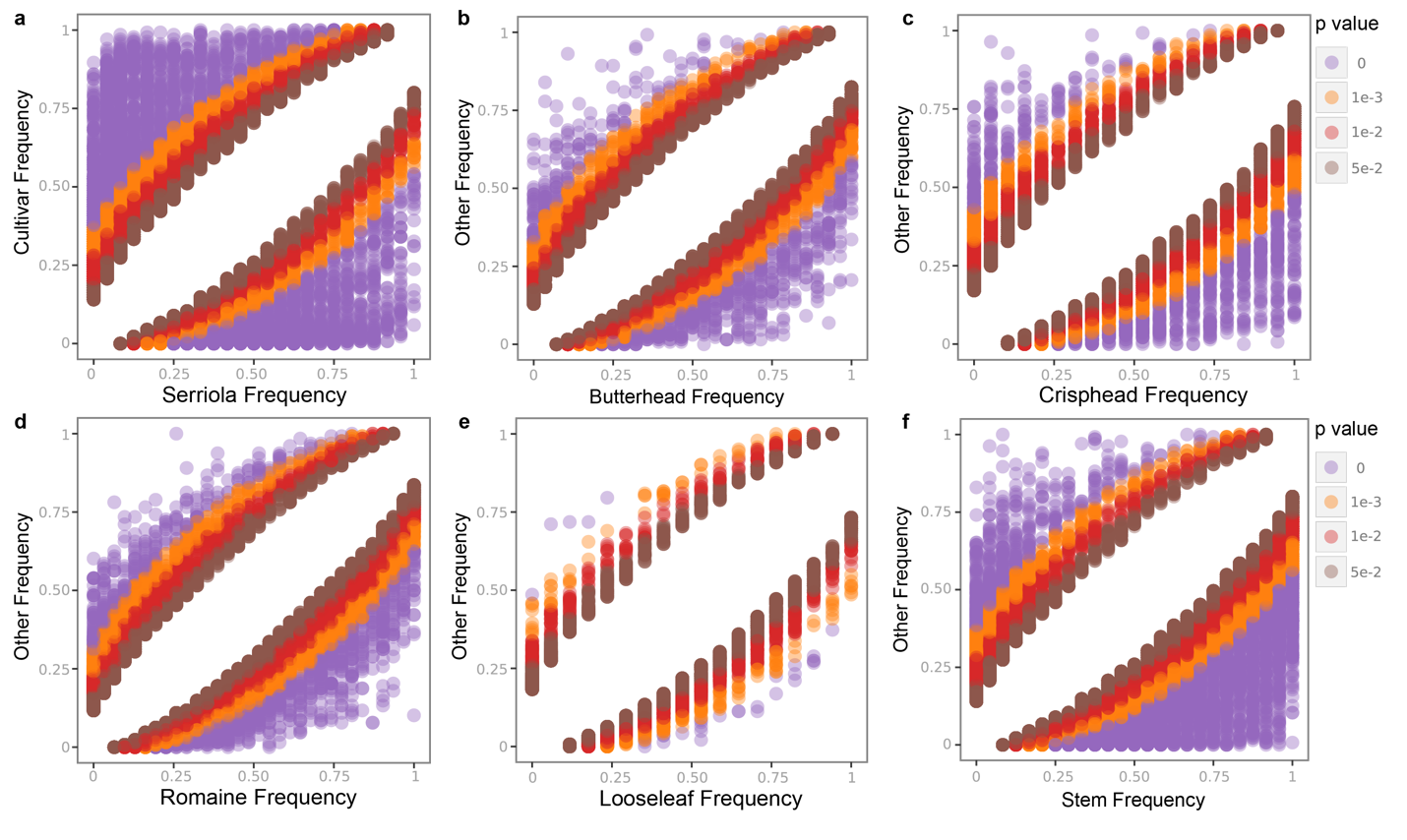

**Supplementary Figure 2. Isoform frequencies during domestication and horticultural diversification**. Scatter plots of isoform occurrence frequency in each focal group versus all other lettuce for a, *L. serriola*, b, Butterhead, c, Crisphead, d, Romaine and e, Looseleaf. f, Stem. Each point represents an isoform; the x-axis shows frequency in the focal group and the y-axis shows adjusted P-value.

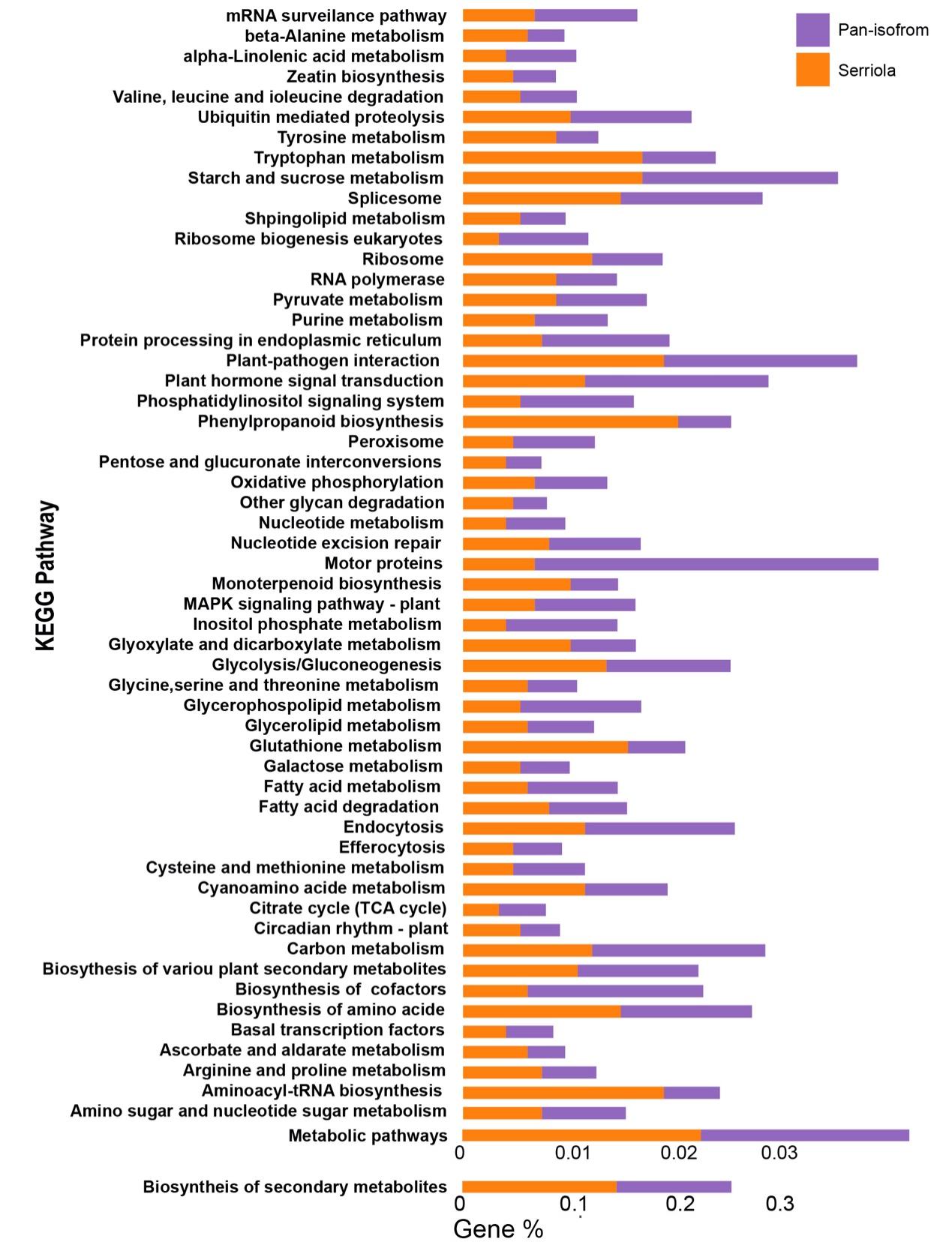

**Supplementary Figure 3. Comparison of KEGG terms annotation between isoforms from *L. serriola* and pan-isoform atlas.** Bars show percentage of genes assigned to each KEGG term, with *L. serriola* and pan-isoform atlas show in orange and purple based on EggNOG annotation terms.

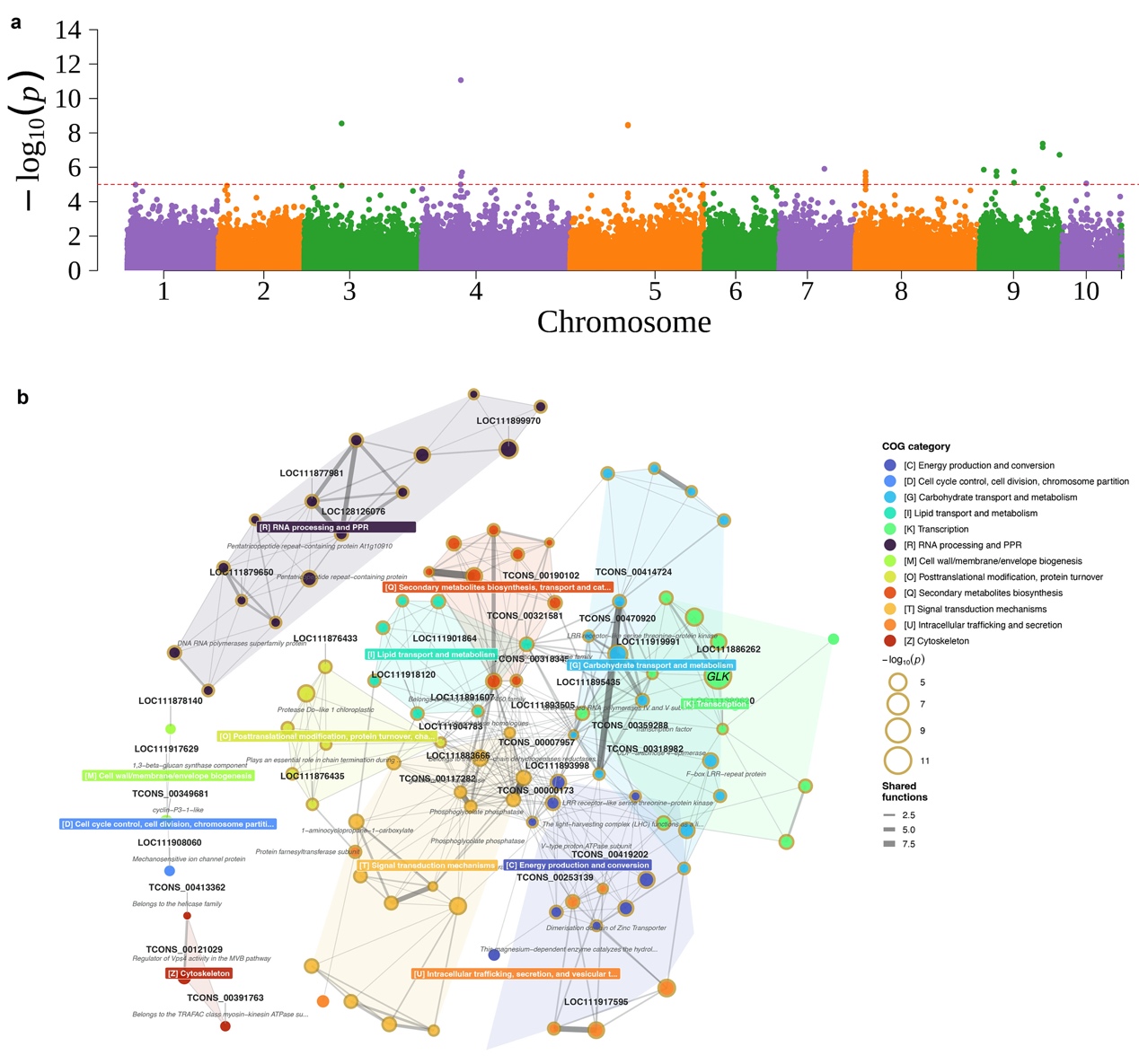

**Supplementary Figure 4. ISONET-GWAS identifies isoforms associated with total leaf pigment and reconstructs co-functional network. a,** Manhattan plot showing the association significance (-log10(p), y-axis) of transcript isoforms across lettuce chromosomes (x-axis). The horizontal dashed red line denotes the genome-wide significance threshold. **b,** Co-functional interaction network of significant candidate isoforms. Nodes represent isoforms and are colored according to their assigned COG functional category, including transcription, secondary metabolism, carbohydrate transport. Node size reflect -log10(p) values, and edge weight represents shared functional scores between connected nodes.

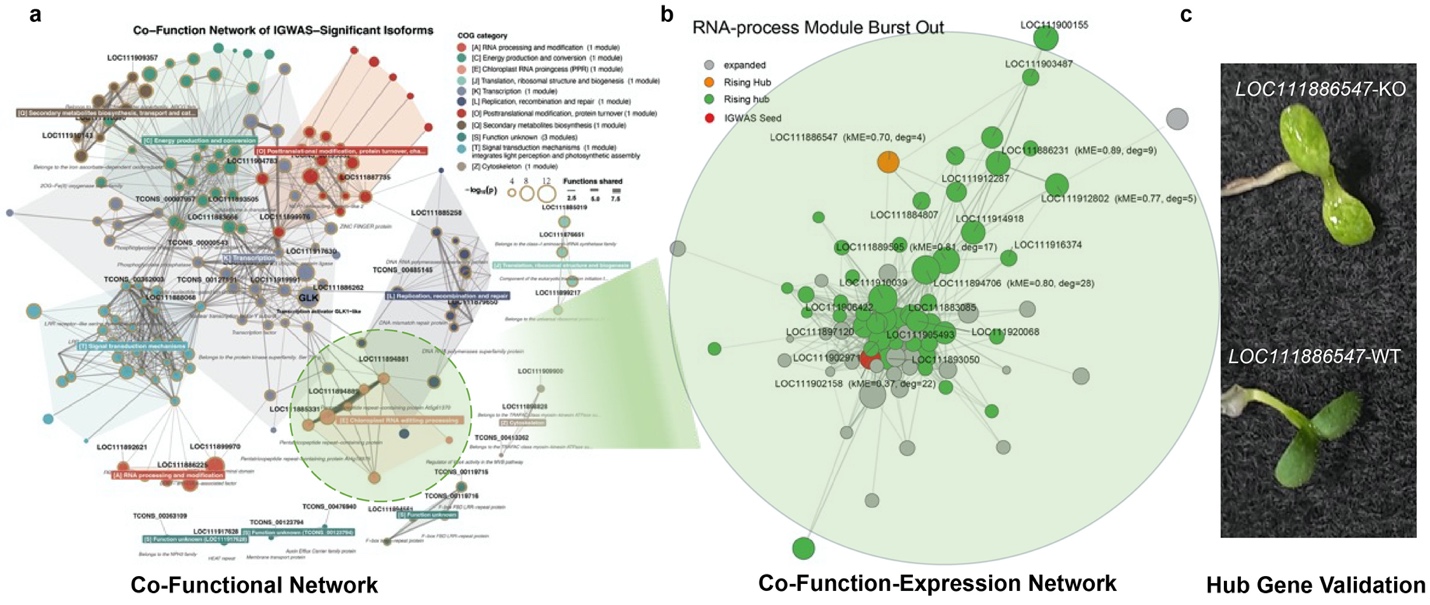

**Supplementary Figure 5 Expansion of co-functional network identified a chloroplast PPR modules associated with chlorophyII regulation**. **a,** Co-functional isoform network of chlorophyll a+b identified using ISONET-GWAS. **b**, Expanded co-functional network of RNA-process module based on co-expression matrix. Module eigengene connectivity (kME) and degree were used to estimate the roles of genes in co-expression network. **c,** Phenotypes of knockout mutants of the candidate hub regulator *LOC111886547* in the co-function-expression network.

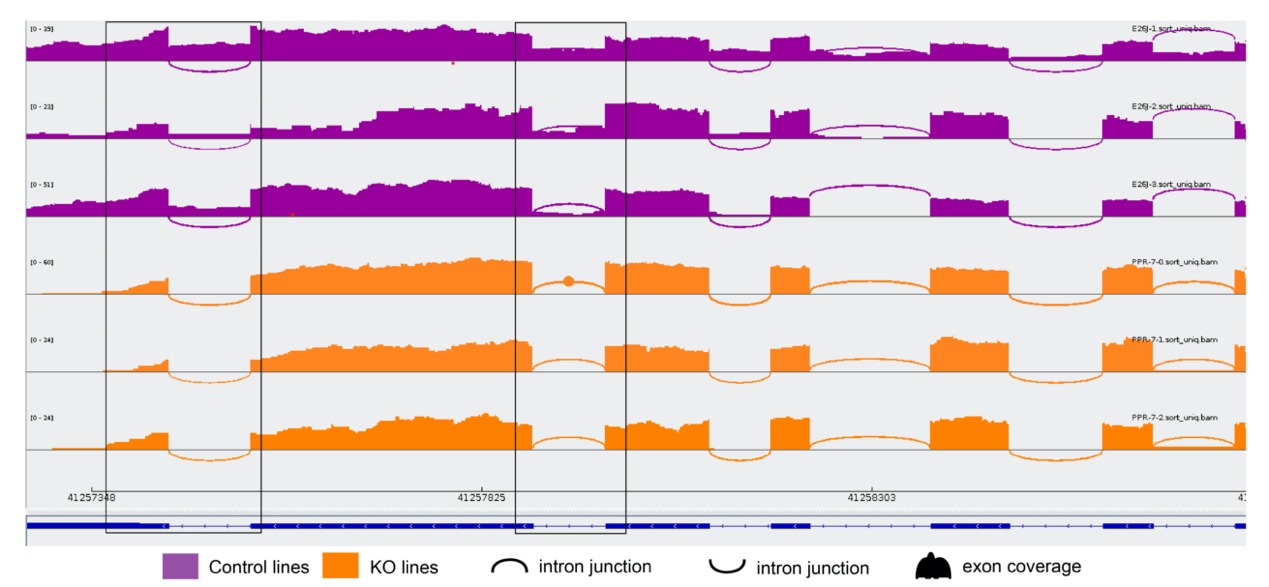

**Supplementary Figure 6. Alternative splicing of *LsFBP24* in *L*s*PPR*-KO lines.** Sashimi plots showing RNA-seq read coverage and splice junction reads across the MADS-box gene *LsFBP24* locus in control and *LsPPR*-KO plants. Tracks are color-coded by genotype, with control lines shown in purple (n = 3) and *LsPPR*-KO lines shown in orange (n = 3). Black outlined boxes highlight specific regions with altered exon coverage or splice-junction usage in *LsPPR*-KO lines relative to control lines.
